# Plasma cell heterogeneity is driven by type of immune challenge

**DOI:** 10.1101/2024.03.26.586773

**Authors:** Mélanie Khamyath, Emilie Lereclus, Vanessa Gourhand, Pierre-Edouard Debureaux, Clémentine Moulin, Hélène Gary, Françoise Bachelerie, Karl Balabanian, Marion Espéli, Amélie Bonaud

**Affiliations:** Université Paris-Cité, Institut de Recherche Saint-Louis, INSERM U1160, F-75010 Paris, France; OPALE Carnot Institute, The Organization for Partnerships in Leukemia, Hôpital Saint-Louis, Paris, France; US31-UMS3679 -Plateforme PLAIMMO, Institut Paris-Saclay d’Innovation Thérapeutique (IPSIT), INSERM, CNRS, Univ.Paris-Sud, Université Paris-Saclay, Clamart, France; Université Paris-Saclay, Inserm, Inflammation, Microbiome and Immunosurveillance, Orsay, France; Control of the immune response B and lymphoproliferation, CNRS UMR 7276, INSERM UMR 1262, University of Limoges, France

**Author notes:** contributed equally Affiliations (Marion Espéli, Amélie Bonaud).

## Abstract

Plasma cells play an essential role in humoral immunity, but many questions remain regarding the heterogeneity of this population, both in terms of ontogeny and involvement in the immune response. In this work, we have identified 5 subsets of plasma cells in human and mouse lymphoid tissues. These subpopulations were distinguished by differential expression of CD62L, CXCR4, FcγRIIb and CD93. The antigenic context as well as the B cell of origin directed plasma cell differentiation towards specific subtypes that display distinct migratory and survival abilities *in vivo*. Altogether, ours results unveil that plasma cell phenotypic and functional heterogeneity relies on intrinsic imprinting during B cell activation.

## Introduction

Plasma cells (PCs) are key players of the immune response through their ability to produce large quantities of antibodies (Ab). Heterogeneity within the PC compartment has previously been reported notably in terms of maturity/longevity, cytokine production and isotype usage^1–7^. However, the function and origin of these different subsets as well as their relevance in human lymphoid organs and in the frame of different immune challenges are not clear^1,8–10^. Here, we identified and characterized 5 subpopulations of PCs easily detectable by flow cytometry in both human and mouse, in spleen and bone marrow (BM) samples. Moreover, we showed that distinct clusters of PCs are induced by different B cell subsets in response to various immune stimuli.

### Study design

Mass and flow cytometry were performed on mouse spleen and BM. Mice were analysed at steady state or after immunization with Sheep Red Blood Cells (SRBC) or NP-Ficoll. All protocols on animals were approved by the relevant institutional review committees (C2EA-26, Animal Care and Use Committee, Villejuif, France, and Comité d’éthique Paris-Nord N°121, Paris, France). PCs were generated *in vitro*^11^, transferred in CD45.1 congenic mice and analysed by flow cytometry. Human BM, blood and spleen samples were collected as approved (Institutional Review Board Paris-Nord, protocol N°10-038 for BM, Biomedicine Agency, protocol n°PFS22-009 for spleen) and stained for flow cytometry analysis (details in supplemental Methods and Table).

## Results/Discussion

To dissect PC heterogeneity, we first developed a panel composed of 24 Ab for mass cytometry (Supp Table 1). By a SPADE on tSNE unsupervised analysis, we highlighted 10 PC subpopulations in the murine spleen (Figure 1A left). Analysis of each marker revealed that some subsets were phenotypically close, differing by only subtle expression of one or two markers (Figure 1A middle and right). Subsets 6 and 7 were the only ones expressing CD62L and had the highest expression of B220 positing them as an early/immature PC cluster^2^. Subsets 1 and 2 were almost similar being only discriminated by the level of expression of CD19 and B220 and thus suggesting that they were forming a homogeneous group composed of PCs at distinct stage of maturation. These subsets, as subsets 3, 4 and 5, formed a continuum characterized by high expression of Cxcr4 and FcγRIIb (detected with a CD16/CD32 Ab). Subsets 8, 9 and 10 had lower expression of both these markers with subsets 8 and 9 expressing more CD93. Based on these observations, we proposed to cluster mouse splenic PCs in 5 subclusters that we termed A to E (Figure 1A, right). We then devised a simplified 4 Ab panel (CD62L, CD16/32, CXCR4 and CD93) to detect these 5 clusters by classical surface flow cytometry (Figure 1B). At basal level in mice, clusters A and B were the most represented in the spleen (23.87 % and 23.52% of total PC respectively), subpopulations C and E represented around 15% of total PC and subpopulation D represented less than 10% of PC. In the BM, cluster B was the most abundant (24.46%), followed by clusters A and E (16.08% and 17.33% respectively), and D and C representing less than 10% of PC (Figure 1C). We next wondered whether these clusters may be related to PC maturation stage, and we combined our 4 Ab panel with CD19 and B220 as markers of PC maturity (Supp. Figure 1A). The three main stages of maturation were observed in all 5 clusters. However, CD19+B220+ plasmablasts (PB) were mainly found in cluster C in both spleen and BM, in agreement with our mass cytometry data^2,12^ (Figure 1A middle and Supp. Figure 1A-B). Most of the very mature PC (CD19-B220-) were found in CD93+ clusters B and E, raising the possibility that they may encompass long-lived PC. As the three stages of maturation were observed in all clusters, it is unlikely that one cluster is at the origin of all the others. While CXCR4 and FcγRIIb expression were already reported on PCs^13,14^, CD62L expression was until now mostly associated with activated/memory B cells^12,15^. CD93 was previously identified in mouse as a marker of longevity in BM PC^16^ but our results suggest that its expression was observed at all stages of PC maturation.

**Figure 1:**
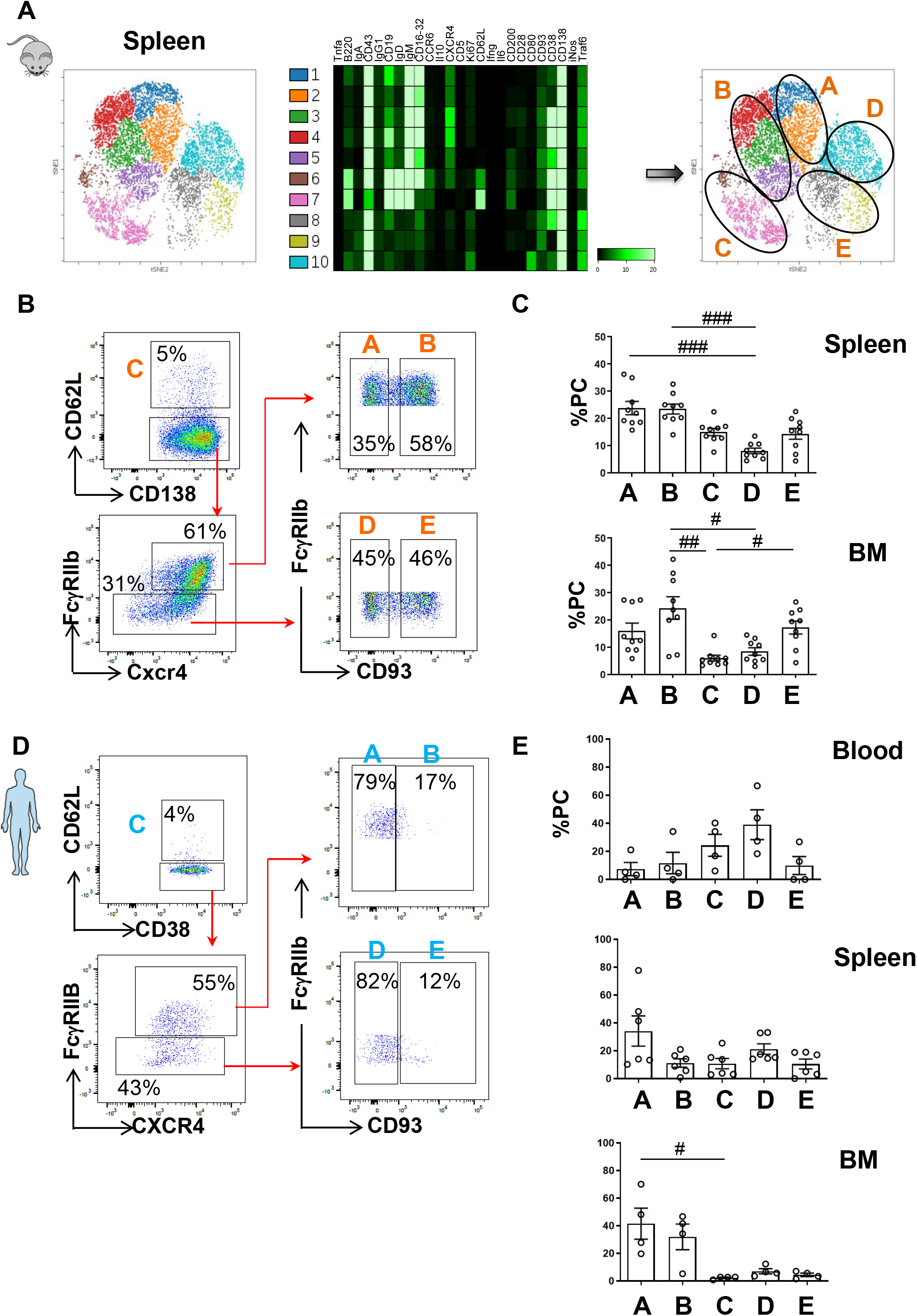
Phenotypic identification of new plasma cell subsets in mouse and human. A) Dimensional reduction representation (viSNE plots) of mass cytometry data of splenic PCs (CD138^+^B220^+/-^) from C57Bl6 mice (left) and corresponding heatmap for protein expression (centre). Subclustering of the 10 subsets in 5 clusters based on protein expression (right). Samples were first normalized and live cells were gated on Ir191/193^+^ and Rh103^-^ cells. One representative experiment shown from 2 independent experiments (n=6 mice). B-E) Gating strategy for mouse (B) and human spleen (D) PC and quantification (C, E) of the 5 subpopulations of PCs in spleen, BM and/or blood identified by flow cytometry. Mouse PC (CD138^+^TACI^+^) and human PC (CD27^+^CD38^+^) subpopulations phenotypes are: A (CD138^+^TACI^+^FcγRIIb^+^Cxcr4^+^), B (CD138^+^TACI^+^FcγRIIb^+^Cxcr4^+^CD93^+^), C (CD138^+^TACI^+^CD62L^+^), D (CD138^+^TACI^+^FcγRIIb^-^Cxcr4^-^), E (CD138^+^TACI^+^FcγRIIb^-^Cxcr4^-^CD93^+^). Cells were gated on their size and structure, on their viability and doublets were excluded. n=9 mice from 3 pooled independent experiments. n=4 for blood and BM sample and 6 for splenic sample from independent human donors. The p-values were determined with a Kruskal-Wallis multiple comparisons test. For ease of reading, only significant differences are shown *ns* > 0.05, *#* p < 0.05, *##* p < 0.01, ### p < 0.001.

We next sought to determine whether a similar gating strategy might be applied to human PC. Strikingly, we found all 5 PC clusters in the human spleen, BM and blood (Figure 1D-E and Supp. Figure 1C). Human spleen was dominated by PC cluster A and cluster D was the second most abundant. The main PC clusters observed in human BM were clusters A and B characterized by a high expression of CXCR4 and FcγRIIB (Figure 1E). In the blood, we observed primarily clusters C and D that were among the least detected in the BM. Altogether, these results unveil the existence of distinct phenotypic clusters of PCs in both mouse and human, easily detectable by flow cytometry and with distinct representation depending on the tissue.

To explore the migratory properties and longevity of each mouse PC subpopulation, we transferred *in vitro* generated PC into recipient mice and analyzed their maintenance for up to 21 days (Figure 2A-C). We first observed that the 5 PC clusters could be generated *in vitro* upon LPS stimulation, with a bias in favor of clusters A, B and C (Figure 2B). One day after transfer, 60% of CD45.2 PC in the spleen and BM were phenotypically part of cluster B (Figure 2C and Supp Figure 2A-B), while they only represented 20% of the transferred cells, suggesting that these cells may have an intrinsic advantage in terms of tissue homing compared to the other PC clusters. Cluster A was the second most abundant in both the spleen and the BM at day 1, while clusters C, D and E were barely detectable in the tissues despite representing a sizable portion of the injected cells (Figure 2C). Interestingly, from day 1 to day 21 we observed a progressive decreased representation of most PC clusters that was mirrored by an increased frequency of cluster E in the spleen and BM (Figure 2C). This pattern may suggest some plasticity among the different subpopulations, with clusters A and B potentially giving rise to cluster E *in vivo*. Alternatively, our results may indicate that cluster E, although migrating less efficiently to lymphoid tissues, is more likely to persist on the long term. This cluster expresses high expression of CD93 that was reported to contribute to long term maintenance of BM PC^16^ although its function in PC biology is not yet completely clear. Our results seem to corroborate this finding, since the 2 main clusters, *i*.*e*. B and E detected at day 21 express this marker. The main difference between these clusters is the low surface expression of Cxcr4 and FcγRIIB in cluster E. The CXCR4/CXCL12 axis has long been described as essential for PC homing to survival niches in both the spleen and the BM^13,17^, potentially explaining the poor migration of cluster E upon transfer. Congruent with our first observations in mouse and human, these results also suggest that cluster C is less likely to be found in lymphoid tissues. Our findings are in line with recent works suggesting that PC longevity determination results from the integration of extrinsic and intrinsic cues and reporting markers for long-lived PC subsets^1,11,18,19^. How these subsets may overlap with the one described here will require further investigation. Taken together, these findings suggest that the PC clusters identified have distinct characteristics and survival ranges.

**Figure 2:**
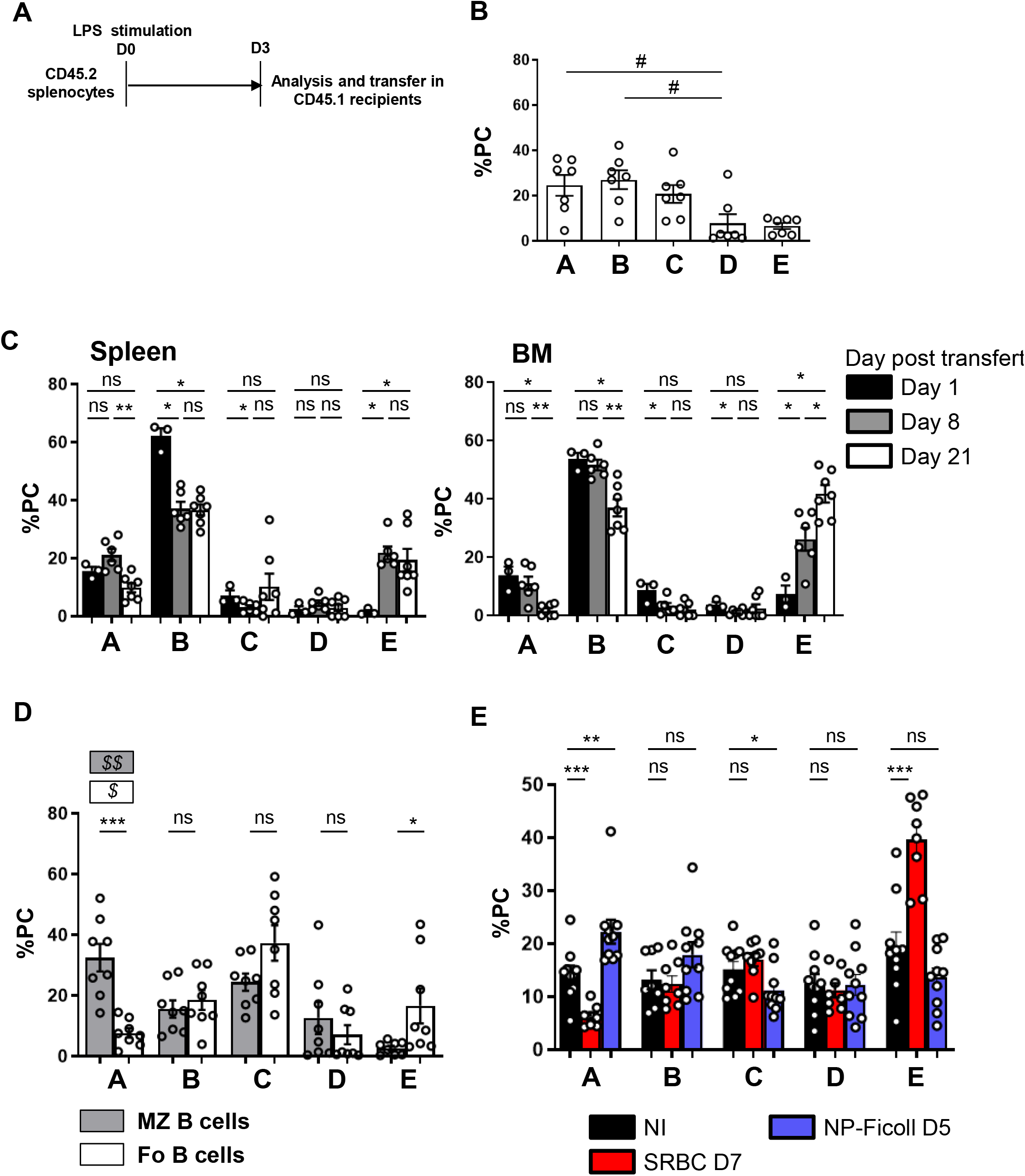
PC clusters have distinct ontogenies and properties. A) Schematic representation of the PC transfer protocol. CD45.2 splenocytes were stimulated during 3 days with LPS and injected intravenously in CD45.1 mice. B) Quantification of PC subpopulations generated *in vitro* from splenocytes after 3 days of stimulation with LPS. C) Frequency of transferred CD45.2 PC subpopulations analysed at day 1, 8 and 21 after transfer, in spleen (left) and BM (right) in CD45.1 mice. n=3-7 mice from 2 pooled independent experiments for each time point. D) Quantification of percentage of *in vitro* generated PC subpopulations after 3 days of LPS stimulation of splenic MZ B cell (grey) or Fo B cell (white). n=8 mice from 3 pooled independent experiments. E) Frequency of splenic PC subpopulations upon distinct immunization or control non-immunized (black). WT mice were immunized with SRBC (red) or NP-Ficoll (blue) and analysed by flow cytometry respectively after 7 days or 5 days of immunization. n=6-8 mice from 2 pooled independent experiments. Cells were first gated on their size and structure, their viability and doublets were excluded. The p-values were determined with the two-tailed Mann-Whitney non-parametric test ns > 0.05, *p < 0.05; ***p < 0.001, and/or a one-way ANOVA test *ns* > 0.05, *$* < 0.05, $$ < 0.01 or a Kruskal-Wallis multiple comparisons test. For ease of reading, only significant differences are shown *ns* > 0.05, *#* p < 0.05, *##* p < 0.01, ### p < 0.001.

We next hypothesized that these distinct PC clusters may have different origins. We thus sorted spleen marginal zone (MZ) and follicular (Fo) B cells and induced PC differentiation *in vitro*. We observed that although MZ and Fo B cells can give rise to all PC clusters to some extent, MZ B cells were the main source of cluster A, while Fo B cells were the main producers of cluster E (Figure 2D and Supp Figure 2C). Next, we assessed whether this was also observed *in vivo*. PC clusters were evaluated after a T-independent immunization with NP-Ficoll stimulating mainly MZ B cells or after a T-dependent immunization with SRBC to stimulate preferentially Fo B cells (Figure 2E). In the context of a T-independent response, we observed a specific increase of PC cluster A mirrored by a decrease of cluster C and a slight reduction of cluster E as detected *in vitro* with sorted MZ B cells. Following SRBC immunization, we observed a strong increase of cluster E associated with a substantial decrease of cluster A, again reflecting what was observed *in vitro* after the stimulation of sorted Fo B cells (Figure 2D-E).

Altogether, our results highlight the emergence of distinct PC subsets depending on the B cell of origin and the type of immune challenge. They extend work by Nutt and colleague that reported a PC specific signature depending on the tissue of origin^20^ and suggest a specific involvement and role for each PC cluster in the course of immune responses. In terms of vaccine development, our findings suggest that a close attention should be paid to the type of PC cluster induced. Moreover, our Ab panel may allow analysis of PC heterogeneity in other conditions including autoimmune diseases and malignant transformation, in which PC play a driver role. Characterisation of their heterogeneity could aid diagnosis and, in the longer term, allow adaptation of therapies according to the PC subpopulations involved.

## Acknowledgements

We thank Dr. N. Setterblad, C. Doliger and S. Duchez (Plateforme technologique IRSL, Paris, France), Dr. V. Parietti (Mouse facility IRSL, Paris, France). The study was supported by ANR @RAction grant (ANR-14-ACHN-0008), ANR JCJC grant (ANR-19-CE15-0019-01) and a grant from IdEx Université Paris-Cité (ANR-18-IDEX-0001) to ME, by the FHU Prothée for EL and KB and the FRM (Programme Equipe FRM 2022, EQU202203014627) to KB. MK was supported by a PhD fellowship from the French Ministry for education and by a 4th year PhD fellowship from “La Ligue contre le cancer”. P-ED had financial support from ITMO Cancer of Aviesan within the framework of the 2021-2030 Cancer Control Strategy on funds administered by Inserm. CM was supported by a Poste d’Accueil Inserm.

## Authorship Contributions

MK performed experiments and analyzed data. EL, VG, PED, CM, performed experiments. HG performed mass cytometry acquisition. FB contributed to mass cytometry experiment design. KB contributed to the project design and to the manuscript redaction. ME designed the project, found funding for the study and wrote the manuscript. AB designed and performed experiments, analyzed data, and wrote the manuscript. All authors had the opportunity to review and edit the manuscript.

## Figure Legends

**Figure supplementary 1:**
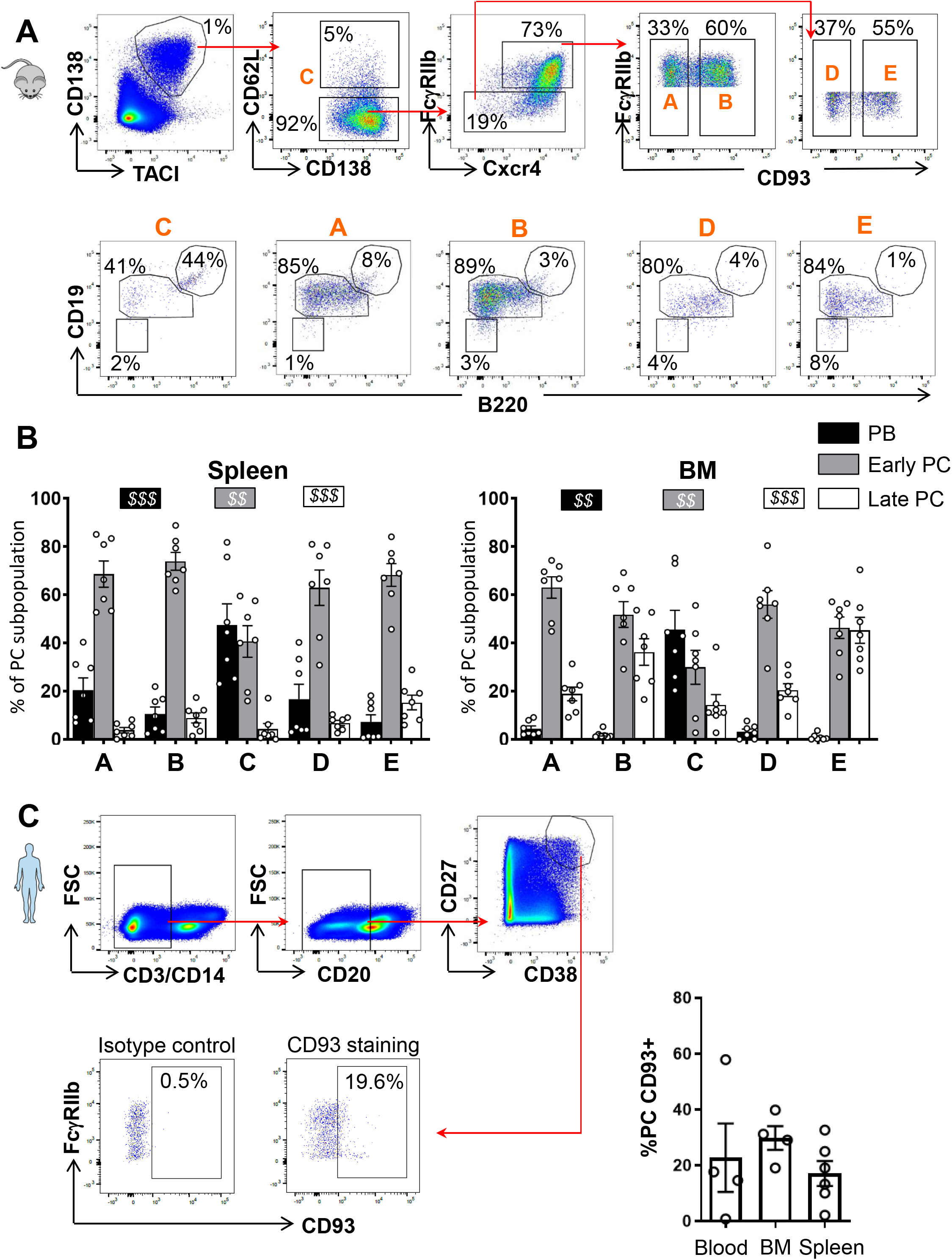
Characterisation of mice PC maturity. A-B) Gating strategy (A) and quantification (B) of the frequency of each PC subset: PBs (CD138^+^TACI^+^B220^+^CD19^+^) (black), early PCs (CD138^+^TACI^+^B220^low^CD19^+^) (grey) and late PCs (CD138^+^TACI^+^B220^-^ CD19^-^) (white) in spleen (B left) and bone marrow (B right) of C57Bl/6 mice. n=7 mice from 3 pooled independent experiments for each time point. C) Gating strategy (left) and quantification (right) of PC (CD3^-^CD14^-^CD20^-/low^) expressing CD93, in human blood, BM and spleen. Cells were first gated on their size and structure, their viability and doublets were excluded. The p-values were determined with a one-way ANOVA test *$$* p < 0.01, *$$$* p < 0.001.

**Figure supplementary 2:**
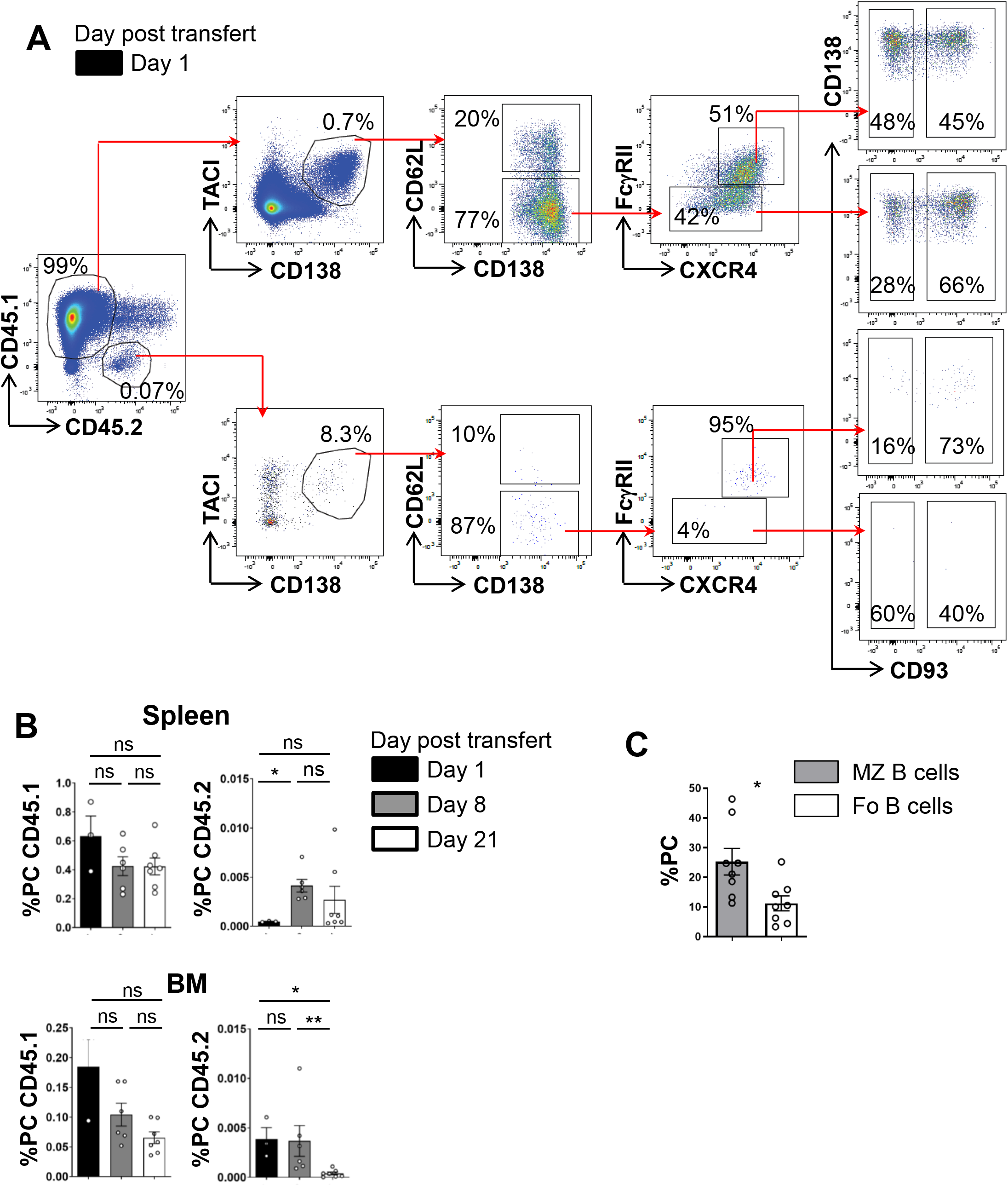
Evolution of PC subpopulations generated in vitro. A-B) Gating strategy of PC subpopulations (day 1) (A) and quantification of CD45.1 and CD45.2 PC on live cells (B) in spleen and BM at day 1, 8 and 21 after intravenous transfer of *in vitro* generated PB. C) Quantification of percentage of in vitro generated PB after 3 days of LPS stimulation of splenic MZ B cell (grey) or Follicular B cell (white). Cells were first gated on their size and structure, their viability and doublets were excluded. n=7-8 mice from 3 pooled independent experiments for each time point. The p-values were determined with the two-tailed Mann-Whitney non-parametric test ns: p > 0.05.

