## Supplementary material for "Plasma cell heterogeneity is driven by type of immune challenge": Supp materials and methods

**Supplemental informations**

**Material and methods**

**Mouse model, immunisation**

All mice were bred in our animal facility under a 12h light/dark cycle in specific pathogen-free conditions (EOPS status). CD57Bl/6J CD45.2 mice (Charles River) aged 8 to 16 weeks were used in all experiments. CD45.1 mice (Charles River) were used as receiver mice in adoptive transfer experiments. All experiments were conducted in compliance with the European Union guide for the care and use of laboratory animals and have been reviewed and approved by appropriate institutional review committees (C2EA-26, Animal Care and Use Committee, Villejuif, France, and Comité d'éthique Paris-Nord N°121, Paris, France). Immunizations were performed intraperitoneally (ip) with 25µg of NP-Ficoll (Biosearch Technologies) or with 200µL of Sheep Red Blood Cells (SRBC) (Eurobio). Daily observation was performed to ensure that no animal was left in a state of pain or suffering during experimentation.

**Human sample preparation**

***Peripheral Blood Mononuclear Cells***

Blood from 4 healthy donor individuals was collected through the local blood bank facility (Etablissement Français du Sang of Saint Louis Hospital, Paris, France). Peripheral Blood Mononuclear Cells (PBMC) were obtained from these donor's cytapheresis by using a ficoll-based density method and stored in fetal calf serum (FCS) supplemented with 10% of DMSO in liquid nitrogen to be further analyzed.

***Spleen***

Cerebral-deceased patients eligible for multi-organ harvesting were included at Saint Louis Hospital (Paris, France) in the protocol n°PFS22-009 approved by the Biomedicine Agency. Spleen samples were collected from 6 patients and stored in PBS at 4°C until they were prepared. Pieces of spleen were first rinsed with PBS, crushed onto a 75µm strainer and finally washed with Phosphate Buffered Saline (PBS). Then, purifying cells was done by using a ficoll-based gradient density method. The cells were stored in FCS supplemented with 10% of DMSO in liquid nitrogen pending analysis.

***Bone Marrow***

Bone marrow samples were obtained during primary total hip replacement surgeries. 4 patients from Orthopedic and Trauma surgery Department of Lariboisiere Hospital (Paris, France) provided written informed consent. Approval from the Institutional Review Board Paris-Nord in the protocol n°10-038 was obtained before the use of the clinical materials for research purposes. Bone fragments were collected in RPMI and stored at 4°C until they were prepared. Bone fragments were washed with PBS, vigorously shaken and allowed to settle, before to collect supernatant into a new tube. This process was repeated several times until bone fragments were white. Pooled supernatants were filtered onto a 100µm strainer and washed with PBS. Finally, cells were purified by using a ficoll-based gradient density method. The cells were stored in FCS supplemented with 10% of DMSO in liquid nitrogen before use.

**Sample isolation in mice**

Spleen and BM cells were isolated from CD57Bl/6J mouse. Medullary cells were obtained after centrifugation of tibias, femurs and hips. Spleens were crushed with a needle piston. Red cells were lysed with ACK for 5min and washed with PBS. Cells were filtered onto a 70µm strainer and resuspended in PBS supplemented with 2% FCS.

### Mass cytometry

5 million cells were collected and stained, firstly with Rh103 for 15min at 37°C. After classical wash (300g-5min), cells were stained with Cisplatin 195Pt for 5min at ambient temperature and washed. Cells were stained with appropriate antibodies (Supplementary Table 1) for cell surface staining during 45min at 4°C. Cells were then washed with MaxParCell buffer (Standard BioTools Inc. ex Fluidigm). Secondary staining with appropriate antibody against fluorochrome (Supplementary Table 1) was performed for 45min at 4°C and washed with MaxParCell buffer (Standard BioTools Inc. ex Fluidigm). Cells were then barcoded as recommended by the supplier (20-Plex PD Barcoding, Standard BioTools Inc. ex Fluidigm). Intracellular staining was then performed with appropriate antibody (Supplementary Table 1) in MaxPar Fix and Perm buffer (Standard BioTools Inc. ex Fluidigm), cells washed and resuspended in MaxParCell buffer in presence of Iridium191/193 (diluted 1/500) until acquisition on Helios (Standard BioTools Inc. ex Fluidigm).

### Flow Cytometry and cell sorting

Murine single-cell suspensions were stained with appropriate antibodies (Supplementary Table 2) in PBS supplemented with 2% Bovine Serum Albumin (BSA) for 30min at 4°C in presence of fixable viability dye eFluor 506 (eBioscience). Cell-sorting experiments were performed using a BD FACS ARIA III cell sorter. MZ and FO cells were harvested in complete culture medium.

Human PBMC purified from blood, spleen and bone marrow were thawed in a 37°C bath and immediately washed with RPMI supplemented with 10% of FCS. Cells were then treated with a 10mg/mL solution of DesoxyriboNuclease I from bovine pancreas (Sigma Aldrich) 15 minutes at 37°C and washed in the same medium. Supernatants were discarded, and cells were resuspended in PBS for cell counting. 5 million cells were collected for staining. Cells were incubated in PBS for 30 minutes at 4°C with the fixable viability dye eFluor 506 (eBioscience) and the antibodies (Supplementary Table 3). After incubation, cells were washed with PBS and resuspended in 300µL of PBS with 1% of paraformaldehyde.

All acquisitions were made using a LSR Fortessa (Becton Dickinson) and conventional analyses were done using FlowJo v10 Software (Becton Disckinson).

### In vitro stimulation

$1x10^6$ splenocytes/mL or $5x10^5$ sorted B cells/mL were stimulated with 1µg/mL of LPS (Invivogen) for 3 days in complete culture medium (RPMI with 10% fetal calf serum, 1% penicillin/streptomycin, 50µM β-mercaptoethanol, 1 mM sodium pyruvate; Gibco, 1mM sodium pyruvate and 0.1 mM nonessential amino acids) with 1U/mL interleukin-4 (IL-4; Miltenyi) and 5 ng/mL IL-5 (Miltenyi).

**Adoptive transfer experiments**

2 x $10^6$ of *in vitro* generated PC (3 days upon LPS+Il4+Il5 as described in section "*in vitro* stimulation") $CD45.2^+$ were transferred intravenously into non irradiated $CD45.1^+$ WT recipients mice. Spleen and BM were harvested and analysed at 1, 8 and 21 days after transfer by flow cytometry.

**Statistical analysis**

The p-values were determined as indicated in the figure legends using the Prism GraphPad software with the two-tailed unpaired Mann-Whitney non-parametric test (*p < 0.05; **p<0.01; ***p<0.001; ****p<0.0001, "ns" = non-significant p-value), or with a one-way ANOVA test for multiple comparisons (*ns* > 0.05, *$* < 0.05, $$ < 0.01) or with a Kruskal-Wallis multiple comparisons test *(ns* > 0.05, # p < 0.05, ## p < 0.01, ### p < 0.001).

**Supplementary Table 1:** List of antibodies used for murin mass cytometry staining

| **Target** | **Tag** | **Clone** | **Company** |
|---|---|---|---|
| CD19 | 149Sm | 6D5 | Standard BioTools Inc. |
| B220 | 144Nd | RA3-6B2 | Standard BioTools Inc. ex Fluidigm |
| CD16/32 | 153Eu | 93 | Standard BioTools Inc. ex Fluidigm |
| CXCR4 | 159Tb | L276F12 | Standard BioTools Inc. ex Fluidigm |
| CD200 | Er168 | OX-90 | Biolegend |
| CD43 | 146Nd | S11 | Standard BioTools Inc. ex Fluidigm |
| CD80 | 171Yb | 16-10A1 | Standard BioTools Inc. ex Fluidigm |
| Traf6 | 173Yb | EP591Y | Abcam |
| iNOS | 161Dy | CXNFT | Standard BioTools Inc. ex Fluidigm |
| CD38 | 175Lu | 90 | Standard BioTools Inc. ex Fluidigm |
| Il6 | 167Er | MP5-20F3 | Standard BioTools Inc. ex Fluidigm |
| IFNg | 165Ho | XMG1.2 | Standard BioTools Inc. ex Fluidigm |
| CD62L | 164Dy | MEL-14 | Standard BioTools Inc. ex Fluidigm |
| Ki67 | 162Dy | B56 | Standard BioTools Inc. ex Fluidigm |
| CD5 | 160Gd | 53-7.3 | Standard BioTools Inc. ex Fluidigm |
| Il10 | 158Gd | JES5-16E3 | Standard BioTools Inc. ex Fluidigm |
| CCR6 | 156Gd | 29-2L17 | Standard BioTools Inc. ex Fluidigm |
| IgM | 151Eu | RMM-1 | Standard BioTools Inc. ex Fluidigm |
| IgG1 | 147Sm | A85-1 | BD Pharmingen |
| TNFa | 141Pr | MP6-XT22 | Standard BioTools Inc. ex Fluidigm |
| IgD | 150Nd | 11-26c-2a | Standard BioTools Inc. ex Fluidigm |
| CD138 | APC | 281-2 | BD Pharmingen |
| APC | 176Yb | APC003 | Standard BioTools Inc. ex Fluidigm |
| IgA | PE | 11-44-2 | Southern Biotech |
| PE | 145Nd | PE001 | Standard BioTools Inc. ex Fluidigm |
| CD93 | FITC | AA4.1 | eBioscience |
| FITC | 174Yb | FIT-22 | Standard BioTools Inc. ex Fluidigm |
| CD28 | Biotin | 37.51 | BD Pharmingen |
| Biotin | 170Er | 1D4-C5 | Standard BioTools Inc. ex Fluidigm |

Antibodies against IgG1, Traf6 and CD200 are manually coupled as recommended par Fluidigm.

**Supplementary Table 2:** List of antibodies used for murin flow cytometry staining

| Target | Fluorochrome | Clone | Company | Dilution |
|---|---|---|---|---|
| B220 | PE-Cy7 | RA3-6B2 | Biolegend | 1/300 |
| CD93 | BV650 | AA4.1 | Becton Dickinson | 1/100 |
| CD21 | PE | 7G6 | Becton Dickinson | 1/900 |
| CD23 | BB700 | B3B4 | Becton Dickinson | 1/200 |
| Live/dead | eFluor506 | | eBioscience | 1/300 |
| CD45.1 | Alexa Fluor 700 | A20 | Biolegend | 1/300 |
| CD45.2 | PE-Cy7 | 104 | Biolegend | 1/300 |
| CD138 | BB700 | 281-2 | Becton Dickinson | 1/300 |
| CD16/32 | APC-Cy7 | 93 | SONY | 1/200 |
| CD200 | BV786 | OX-90 | Becton Dickinson | 1/100 |
| CXCR4 | PE | 2B11/CXCR4 | Becton Dickinson | 1/50 |
| CD62L | BV421 | MEL-14 | Becton Dickinson | 1/50 |
| CD38 | BV605 | 90/CD38 | Becton Dickinson | 1/100 |
| TACI | APC | 8F10-3 | eBioscience | 1/100 |
| B220 | FITC | RA3-6B2 | Becton Dickinson | 1/300 |
| CD19 | PE-CF594 | 1D3 | Becton Dickinson | 1/500 |

**Supplementary Table 3:** List of antibodies used for human flow cytometry staining

| Target | Fluorochrome | Clone | Company | Dilution |
|---|---|---|---|---|
| CD20 | PerCP-Cy5.5 | 2H7 | Becton Dickinson | 1/100 |
| CD19 | FITC | HIB19 | BioLegend | 1/100 |
| CD38 | BV786 | HIT2 | Becton Dickinson | 1/100 |
| CD27 | BV650 | L128 | Becton Dickinson | 1/50 |
| CD93 | APC | REA1111 | Miltenyi | 1/50 |
| CXCR4 | BUV737 | I2G5 | Becton Dickinson | 1/100 |
| CD3+CD14 | V450 | UCHT1/MΦP9 | Becton Dickinson | 1/100 |
| CD62L | PeCF594 | DREG-56 | Becton Dickinson | 1/50 |
| CD32 | PE | FLI8.26 | Becton Dickinson | 1/100 |
| CD138 | BV605 | MI15 | Becton Dickinson | 1/100 |
